# CryoET reveals actin filaments within platelet microtubules

**DOI:** 10.1101/2023.11.24.568450

**Authors:** Chisato Tsuji, Marston Bradshaw, Megan Allen, Molly L. Jackson, Judith Mantell, Ufuk Borucu, Alastair W. Poole, Paul Verkade, Ingeborg Hers, Danielle M. Paul, Mark P. Dodding

## Abstract

Crosstalk between the actin and microtubule cytoskeletons is essential for many cellular processes. Recent studies have shown that microtubules and F-actin can assemble to form a composite structure where F-actin occupies the microtubule lumen. Whether these cytoskeletal hybrids exist in physiological settings and how they are formed is unclear. Here, we show that the short-crossover Class I actin filament previously identified inside microtubules in human HAP1 cells is cofilin-bound F-actin. Lumenal F-actin can be reconstituted *in vitro*, but cofilin is not essential. Moreover, actin filaments with both cofilin-bound and canonical morphologies reside within human platelet microtubules under physiological conditions. We propose that stress placed upon the microtubule network during motor-driven microtubule looping and sliding may facilitate the incorporation of actin into microtubules.

## Introduction

Crosstalk between the actin and microtubule cytoskeletons is essential for many cellular processes [1, 2]. This is thought to be mediated by proteins that directly connect or signal between the two dynamic networks to coordinate their activities. However, recent studies have challenged this framework by showing that microtubules and F-actin can assemble to form an unanticipated composite structure where F-actin occupies the lumen of the cylindrical tubulin polymer [3, 4].

Lumenal actin filaments were identified in human HAP1 cells treated with a small molecule that targets kinesin-1 to induce the formation of thin, membrane bound, microtubule-based projections that are highly accessible to cryo-electron tomography (cryoET) [4]. In this system, microtubules form a dynamic bundle and lumenal actin filaments are highly abundant, with morphologies (named Class I and Class II) that are distinct from the canonical cytoplasmic/muscle form; principally, the cross-over spacing between the two ‘long-pitch’ strands of the actin double helix is short (at around 27 nm), compared to the canonical 35–37 nm [5-7]. This suggested that actin binding proteins (ABPs) which modify the twist of the filament may be present [8, 9].

CryoET analysis of *Drosophila* S2 cells revealed similar filaments in microtubule-based projections in cells treated with the actin-targeting drug cytochalasin D, whose abundance was enhanced by additional treatment of cells with thapsigargin [3]. Sub-tomogram averaging and knockdown studies demonstrated that these filaments are composed of cofilin-bound F-actin (cofilactin); cofilin is known to change the twist of F-actin [9, 10].

Together, these studies show that F-actin can occupy the microtubule lumen in chemically-induced microtubule-based projections emerging from both human and insect cells in culture. However, basic requirements for formation of these structures are unknown. Moreover, the impact of these small-molecule manipulations and the fact that these studies were performed on cultured cells, leaves the physiological occurrence of lumenal actin unclear.

Here, we examine the composition of lumenal F-actin in human HAP1 cells and show that the Class I filament is composed of cofilin-bound F-actin. To explore the requirements for the formation of these structures, we utilise an *in vitro* reconstitution system which demonstrates that, although cofilin-bound F-actin can incorporate into microtubules, cofilin is not essential. A dynamic and bundled microtubule cytoskeleton is present in native human platelets [11] and Focussed-Ion-Beam (FIB) milling and cryoET reveals that actin filaments with both cofilin-bound and canonical morphologies are found within platelet microtubules, providing an unequivocal identification of lumenal F-actin in a physiological setting that is not modified by small-molecule treatment.

## Results and Discussion

The cofilin-bound F-actin recently identified within the microtubule lumen in *Drosophila* S2 cells is morphologically similar to the Class I filament observed in human HAP1 cells [3, 4] There are two shared features of particular note; firstly, the short spacing of the ‘crossovers’ of the actin double helix that can be readily identified and directly measured in high-quality tomograms and are apparent as layer lines in their power spectra [4]; and secondly, the smooth appearance of the filament that is distinct from the classic ‘beads on a string’ appearance of canonical F-actin [12]. To determine whether human Class I filaments are cofilin-bound F-actin, we treated HAP1 cells with the kinesin-1 targeting small-molecule ‘kinesore’ to induce projections using our established protocol and imaged those projections with cryoET using a 300kV Titan Krios microscope **(Fig. 1A-C)** [4, 13]. In this new dataset, we observed lumenal actin filaments corresponding to the Class I and Class II forms **(Fig. 1B and Fig. 1D, top left)** [4]. Class II filaments showed an F-actin-like power spectrum with short crossover spacing augmented with a prominent meridional layer line indicating a deviation from the typical F-actin helical morphology (**Fig. S1A**). We also observed rarer examples of a morphologically distinct filament that we named Class III that did not display an F-actin-like power spectrum **(Fig. 1D, top right, Fig. S1B)**. Nonetheless, images showing transitions between filament classes through breaks or less well-defined intermediates suggest that each of these filaments are likely to be actin structures, perhaps undergoing disassembly or assembly, or with different binding partners **(Fig. 1D, middle and bottom**). This opens the intriguing possibility that lumenal actin is dynamic.

**Fig. 1.**
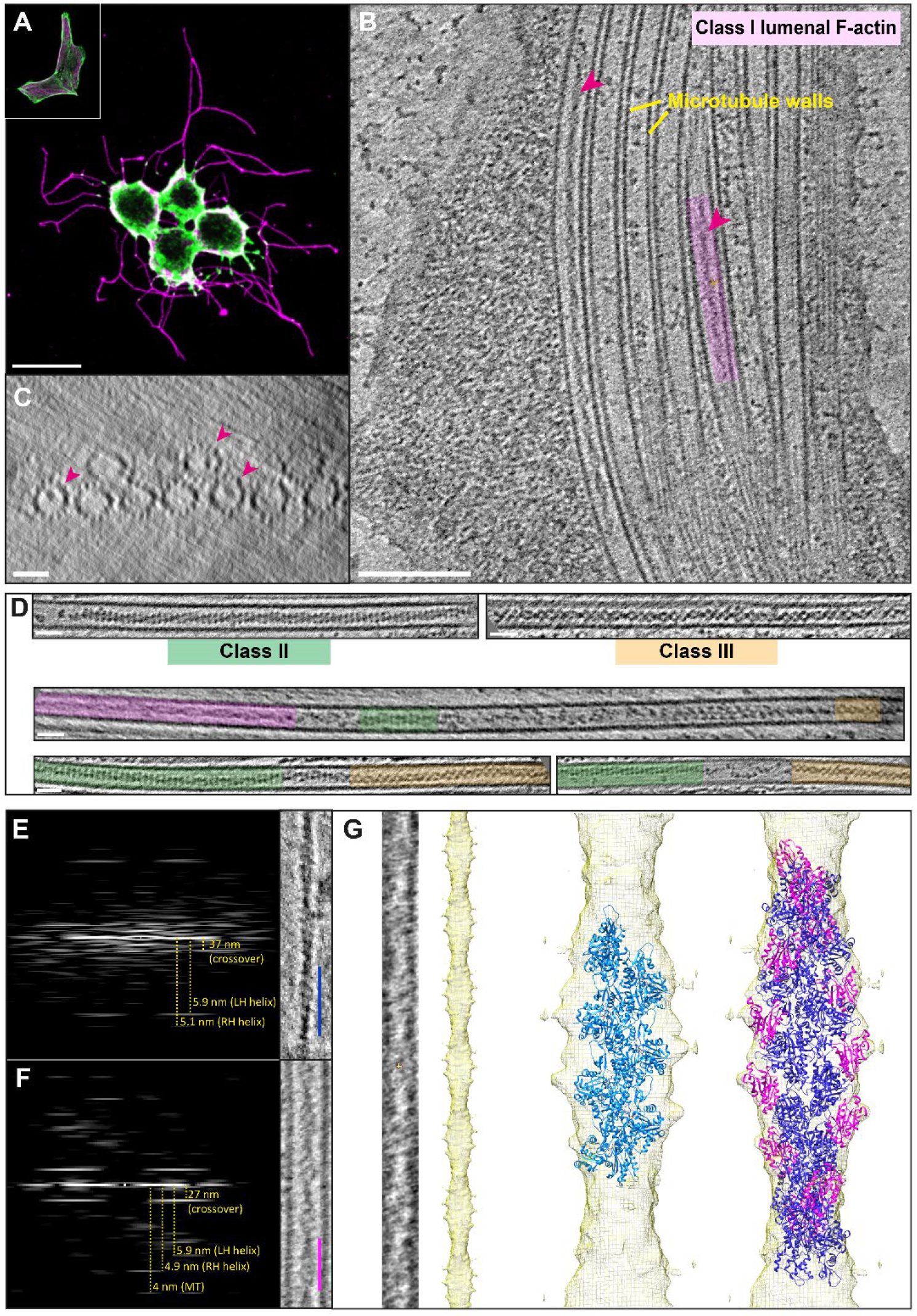
Morphology and composition of lumenal actin filaments. **(A)** Fluorescence microscopy images showing control (inset) and kinesore-treated (main image)) HAP1 cells stained using antibodies targeting beta-tubulin (magenta) and actin (green). Scale bar = 20 μm. **(B**,**C)** Tomogram slices (in longitudinal (B) (Scale bar = 25 nm) and transverse orientations (C) (Scale bar = 100 nm)) showing microtubules within a projection from a kinesore-treated HAP1 cell. Class I lumenal filaments are highlighted with magenta arrows and one example is boxed in magenta. **(D)** Examples of Class II and Class III filaments (top) and transitions between filament morphologies (middle and bottom). Class I is shaded magenta, Class II is green and Class III is orange (Scale bar = 25 nm). **(E)** Layer lines from an *in vitro* example (right, blue scale bar = 37 nm) of a canonical F-actin labelled with real space distances annotated. **(F)** Layer lines of an example class I filament (from HAP1 cells) (right, magenta scale bar = 27 nm) labelled with real space distances. **(G)** An example of a segment of class I filament that has been straightened, inverted, and projected in z, which was averaged to produce a helical reconstruction map. When structures of actin (PDB: 8D17, light blue) was docked in, it was insufficient to fill the map, whereas cofilin-actin (PDB: 3J0S, actin in dark blue, cofilin in magenta) is in good agreement with the model.

Of 51 lumenal filaments in this new dataset, 35 displayed the Class I morphology, with a crossover spacing (measured directly from layer lines) of 27.66 nm (+/-0.23 s.d.) that contrasted with the long cross-over spacing of a canonical actin filament (**Fig. 1E, F**). We performed helical reconstructions to generate 3D maps on the five highest quality images, and one representative example is shown **(Fig. 1G**). Docking of a cryo-electron microscopy (cryoEM) structure of canonical F-actin (light blue) (PDB: 8D17) [14] into the Class I helical reconstruction model was insufficient to explain the density. In contrast, when a cryoEM cofilin-actin structure (dark blue and magenta) (PDB: 3J0S) [15] was docked in, it fitted our model well (cross correlation score 0.92) **(Fig. 1G)**. Similarly, our model fitted well with density of cofilin-actin filaments from the lumen of microtubules in S2 cells **(Fig. S1C)** [3]. Together, these data enable us to conclude that the Class I filament in HAP1 cells is cofilin-bound F-actin.

To define basic requirements for the incorporation of F-actin into microtubules, we used an *in vitro* reconstitution system, taking advantage of the ‘TicTac’ buffer that allows dynamic assembly of both polymers [16] **(Fig. 2)**. When purified actin and tubulin were polymerised together, both F-actin and microtubules could be observed using cryoET **(Fig. 2 A)**. In samples where cofilin was present, the short crossovers and distinct morphology of the actin filaments was readily apparent (pink arrowheads) [9], confirming the proposition that these characteristics are a good proxy for the presence of high levels of bound cofilin in cryoET images of F-actin *in situ* **(Fig. 2B)**. Occasionally, F-actin and cofilin-bound filaments were found to run alongside microtubules, but most formed a lose mesh that was not microtubule associated. In both conditions, we were able to observe lumenal filaments in a minority of the microtubules (≈ 2% for F-actin alone and ≈ 5% for cofilin-actin) **(Fig.2 C,D)**. The appearance of these *in vitro* lumenal cofilin-actin filaments was indistinguishable from Class I filaments observed *in situ* in HAP1 cells. We also noted examples of actin/cofilin-actin filaments apparently associated with breaks in the microtubule lattice or emerging from microtubule ends (**Fig. 2E,F**). Thus, both F-actin and cofilin-bound F-actin can incorporate into the microtubule lumen. Although cofilin-bound F-actin is the predominant form in HAP1 and S2 cells (pointing to a functional or regulatory role for cofilin), addition of cofilin is not essential for F-actin incorporation in this simple *in vitro* system.

**Fig. 2.**
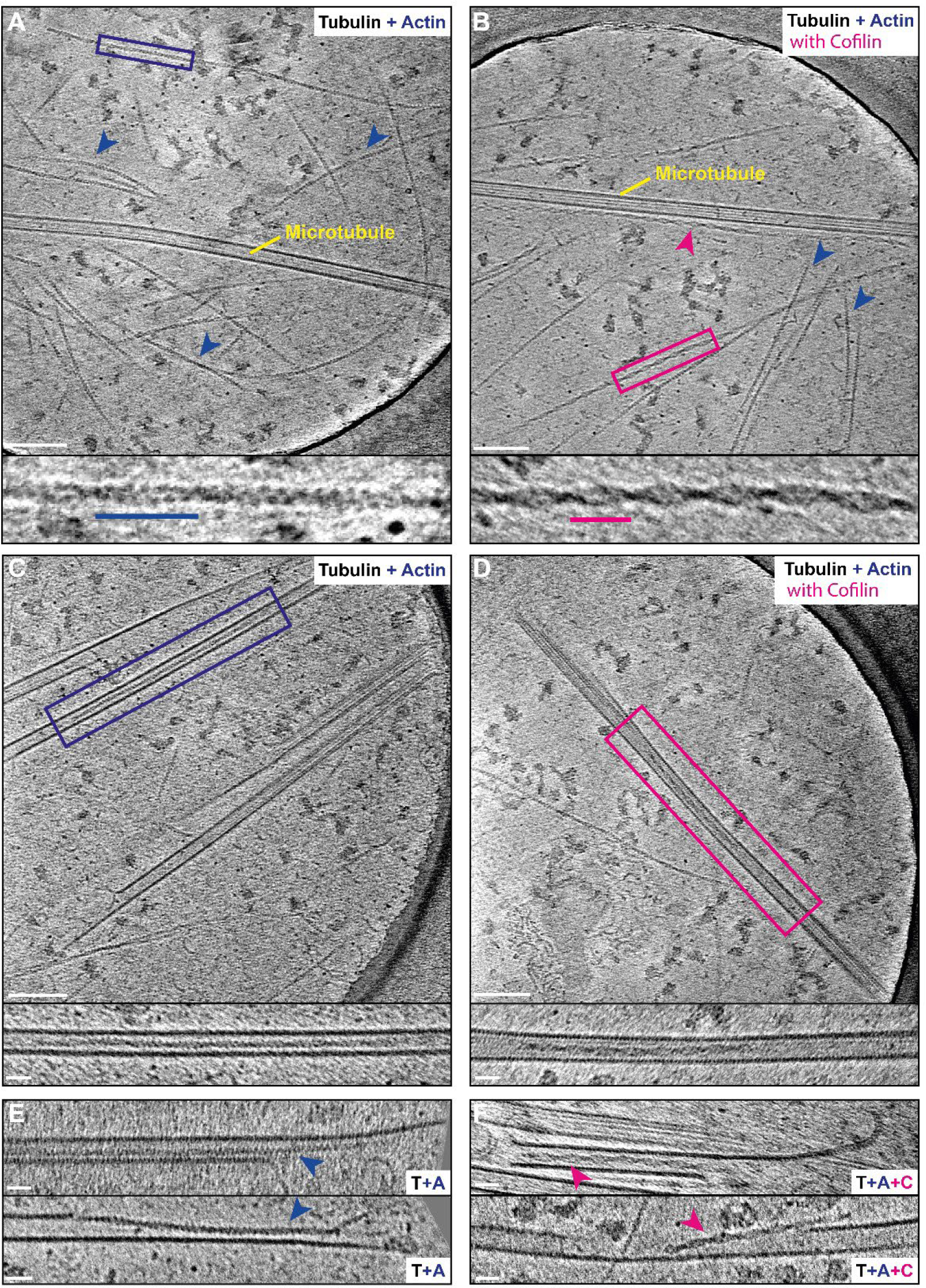
In vitro reconstitution of lumenal F-actin and cofilin-bound F-actin. **(A)** Representative slice from a tomogram showing organisation of microtubules and actin filaments that have been polymerised together. (Scale bar = 100 nm). Expanded box highlights the canonical ‘beads-on-a-string’ F-actin morphology and long crossover spacing (blue line is 37 nm). **(B)** Representative slice from a tomogram showing typical organisation of microtubules and actin filaments that have been polymerised together in the presence of cofilin. Expanded box highlights the distinctive smooth appearance of the cofilin-bound actin filaments and short-crossover spacing (magenta line is 27 nm). **(C)** Representative slice from a tomogram showing a canonical actin filament within a microtubule, boxed and expanded below. **(D)** Representative slice from a tomogram showing a cofilin-bound actin filament within a microtubule, boxed and expanded in below. **(E)** Examples of actin-filaments at microtubule ends or breaks in the lattice. **(F) E**xamples of cofilin-bound actin-filaments at microtubule ends or breaks in the lattice.

We next sought to establish whether lumenal actin filaments exist in native or primary cells under physiological conditions without small-molecule treatments. We considered that one factor in common between systems where microtubule lumenal F-actin has been unambiguously observed is that microtubules are most likely under significant mechanical stress, as the formation of extended microtubule-based projections is driven by microtubule motors [4, 17]. Indeed, projections in HAP1 cells form through the extrusion of tight microtubule loops that push against the plasma membrane [4]. CryoET analysis of the tips of these loops show evidence of extreme microtubule curvature, breakage and depolymerisation and the presence of all three filament classes (both in and outside of the microtubule lumen), **(Fig. S2; Supplementary Movie 1**). Ostensibly similar motor-driven microtubule looping is important for extrusion in platelet biogenesis from mega-karyocytes [18] resulting in a dynamic circumferential bundle of microtubules within platelets themselves. This characteristic bundle, called the marginal band, mechanically maintains the platelet discoid shape during their resting state and rapidly remodels under platelet activation [11, 19-21].

Because of the relative thickness of platelets (compared to small-molecule induced projections), we turned to FIB-milling. Human platelets were put on EM grids, plunge frozen, thinned by FIB-milling and analysed using cryoET **(Fig. 3A)**. We observed bundled microtubule structures which are consistent with the marginal band [22-24] (**Fig. 3B,C; Supplementary Movie 2)**, as well as more isolated microtubules **(Fig. 3D,E)**. Within the microtubules, actin filaments were readily identified in 6 out of 25 tomograms with visible sections of microtubule lumen, from 2 independent platelet preparations. Consistent with previous observations in HAP1 and S2 cells, the majority had the distinctive Class I cofilin-bound F-actin morphology (**Fig. 3B,C,D insets**), although we also observed less frequent examples of long-crossover filaments more closely resembling the canonical cofilin-free form, consistent with our *in vitro* reconstitution assays **(Fig. 3E, inset)**.

**Fig. 3.**
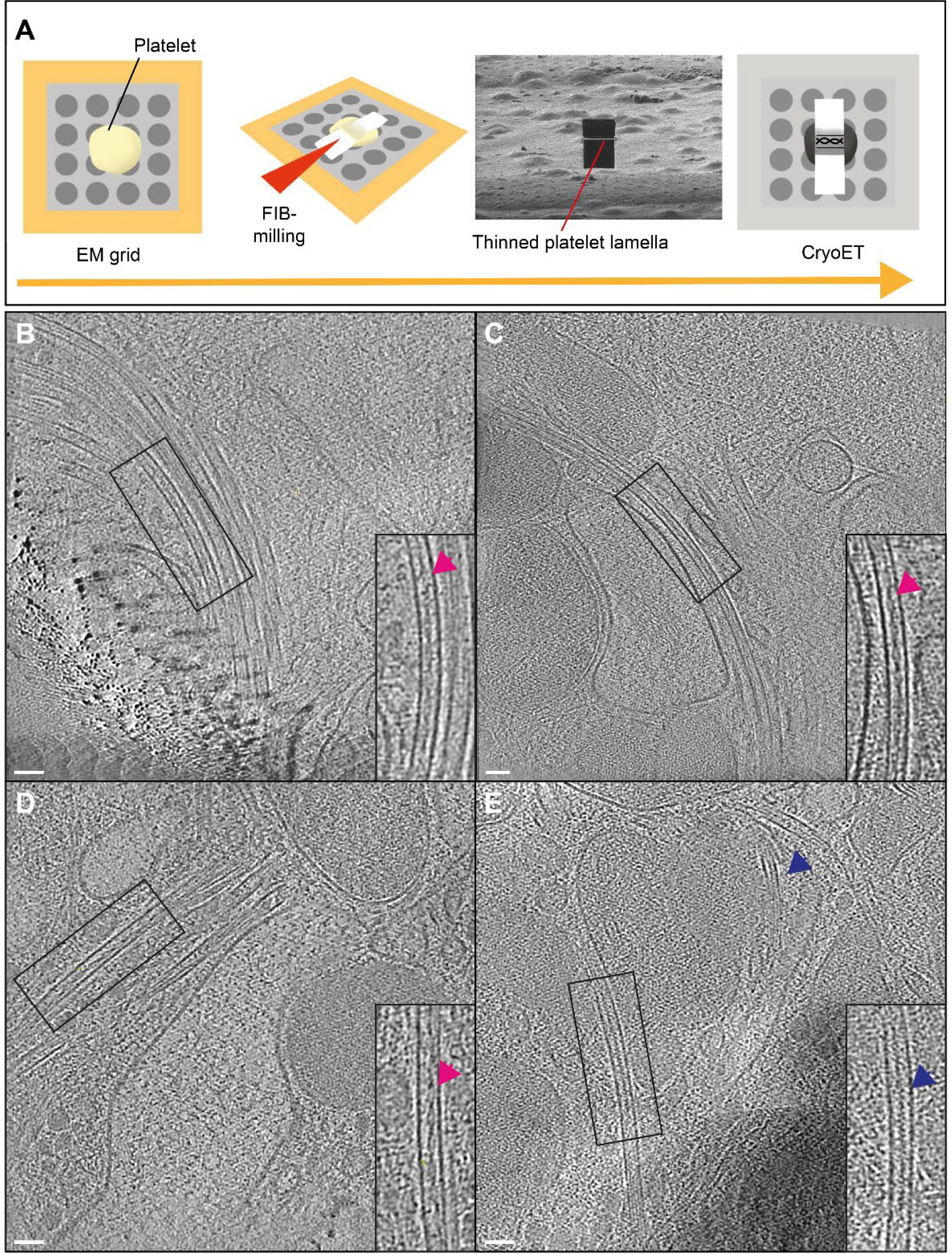
F-actin resides within the lumen of human platelet microtubules. **(A)** Schematic showing plalelet FIB-milling and cryoET workflow. **(B-E)** Representative slices from tomograms showing cofilin-bound actin filament within platelet microtubules (magenta arrows) and an actin filament with a canonical morphology (blue arrow). (Scale bar = 50 nm). (B) Is a 13-microtubule bundle. One microtubule contains an actin filament with Class I morphology, seven appear empty or contain globular densities. Lumenal contents of the remaining microtubules were ambiguous. (C) Is a 11-microtubule bundle. Two microtubules contain filaments with the Class I morphology, seven appear empty or contain globular densities. Contents of the remaining microtubules were ambiguous. A video showing a Z-series though (C) is provided in Supplementary Movie 2.

We did not observe clear examples of Class II and Class III filaments in platelets or in *vitro*. This may reflect the relatively high abundance of lumenal filaments in the HAP1 cell system, or a missing component in the *in vitro* reconstitution system. In any case, the presence of these structures in HAP1 cell projections highlights the importance of considering intermediates on an assembly/disassembly pathway. This may be particularly important when examining morphologically diverse lumenal material in other cell types that has filamentous character but unknown composition [25]. Structural studies of Class II and Class III filaments are therefore a priority.

In summary, we have characterised the form and composition of microtubule lumenal F-actin *in situ*, reconstituted cofilin-free and cofilin-bound forms *in vitro*, and demonstrated that lumenal F-actin is present in *ex vivo* primary human cells without chemical modification. Going forward, it will be useful to explore the effect of mutations in cytoskeletal genes that cause hereditary thrombocytopenias, including in the cofilin pathway, for effects on lumenal F-actin [26-28]. More broadly, this challenge to the textbook representation of cytoskeletal composition and crosstalk should lead to a sustained effort to understand the function and regulation of lumenal actin in platelets and determine its presence in other cell types. These latter efforts should focus on settings where microtubules are under structural and mechanical stress. Our findings suggest that a full understanding will require development of new models of single filament actin dynamics in confined environments and new experimental systems to explore the impact of lumenal F-actin on microtubule properties.

## Author contributions

Conceptualization: CT, PV, DMP, AWP, IH, MPD.

Formal analysis: CT, MB, JM, IH, PV, AWP, DMP, MPD

Investigation: CT, MB, JM, IH, PV, AWP, DMP, MPD

Methodology: CT, MB, MA, MLJ, JM, UB

Writing – original draft: CT, MPD

Writing – reviewing and editing: All authors

## Acknowledgements

C.T. is supported by the Wellcome Trust 4-year PhD Programme in Dynamic Molecular Cell Biology (203959/Z/16/Z) at the University of Bristol. M.B. is supported by a British Heart Foundation on a PhD Studentship FS/20/5/34973. M.A and M.L.J. are supported by the British Heart Foundation 4-year PhD Programme in Integrative Cardiovascular Sciences (FS/4yPhD/F/20/34125). AWP is a Wellcome Trust Investigator (219472/Z/19/Z) and is supported by grants from the British Heart Foundation (SP/F/21/150023; PG/21/10760; FS/19/53/34887). Work in the lab of I.H. is supported by the UKRI Biotechnology and Biosciences Research Council (BB/X017176/1) and the National centre for Replacement, Refinement and Reduction of Animals in Research. D.M.P. has been supported by the British Heart Foundation through a Career Re-Entry Fellowship (FS/14/18/3071) and a current Intermediate Basic Science Research Fellowship FS/IBSRF/23/25156. This work was also supported by a Lister Institute of Preventative Medicine Fellowship to M.P.D and work in his lab is supported by the UKRI Biotechnology and Biosciences Research Council (BB/W005581/1).

We thank Andrew P. Carter and Camilla V. Santos (MRC-LMB) for comments on and support with the project, and Michael Way (Francis Crick Institute) and Edward H. Egelman (University of Virginia) for helpful discussions. We acknowledge access and support of the GW4 Facility for High-Resolution Electron Cryo-Microscopy, funded by the Wellcome Trust (202904/Z/16/Z and 206181/Z/17/Z) and BBSRC (BB/R000484/1). We acknowledge Diamond for access and support of the cryo-EM facilities at the UK national electron Bio-Imaging Centre (eBIC), proposals BI25452-24, BI32707-3, BI32707-7, BI32707-12, and BI32707-14, funded by the Wellcome Trust, MRC, and BBSRC.

## Ethics statement

Approval for the platelet work in this study was granted to IH by South Central—Hampshire A Research Ethics Committee (NHS-REC reference 20/SC/0222).

## Data availability

EMDB and EMPIAR depositions are in progress, and this preprint will be updated with accession numbers when this process is complete.

## Supplementary Figures

**Fig. S1.**
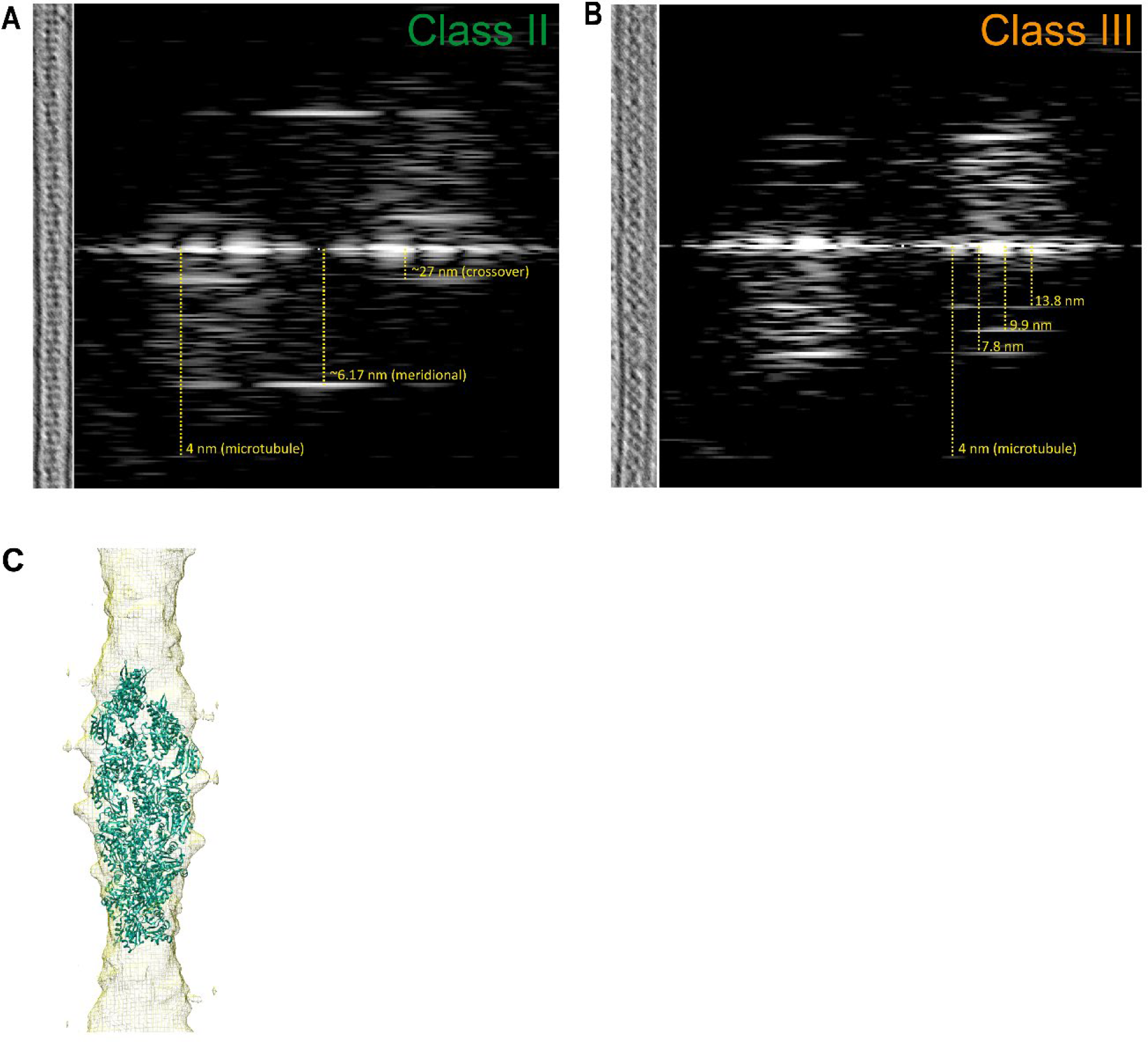
Characterisation of lumenal filaments. (A,B) Power spectra of Class II and Class III filaments annotated with real space distances. (C) Docking of cofilin-actin structure from Ventura Santos *et al. [3]*. into map shown in Fig. 1G.

**Fig. S2.**
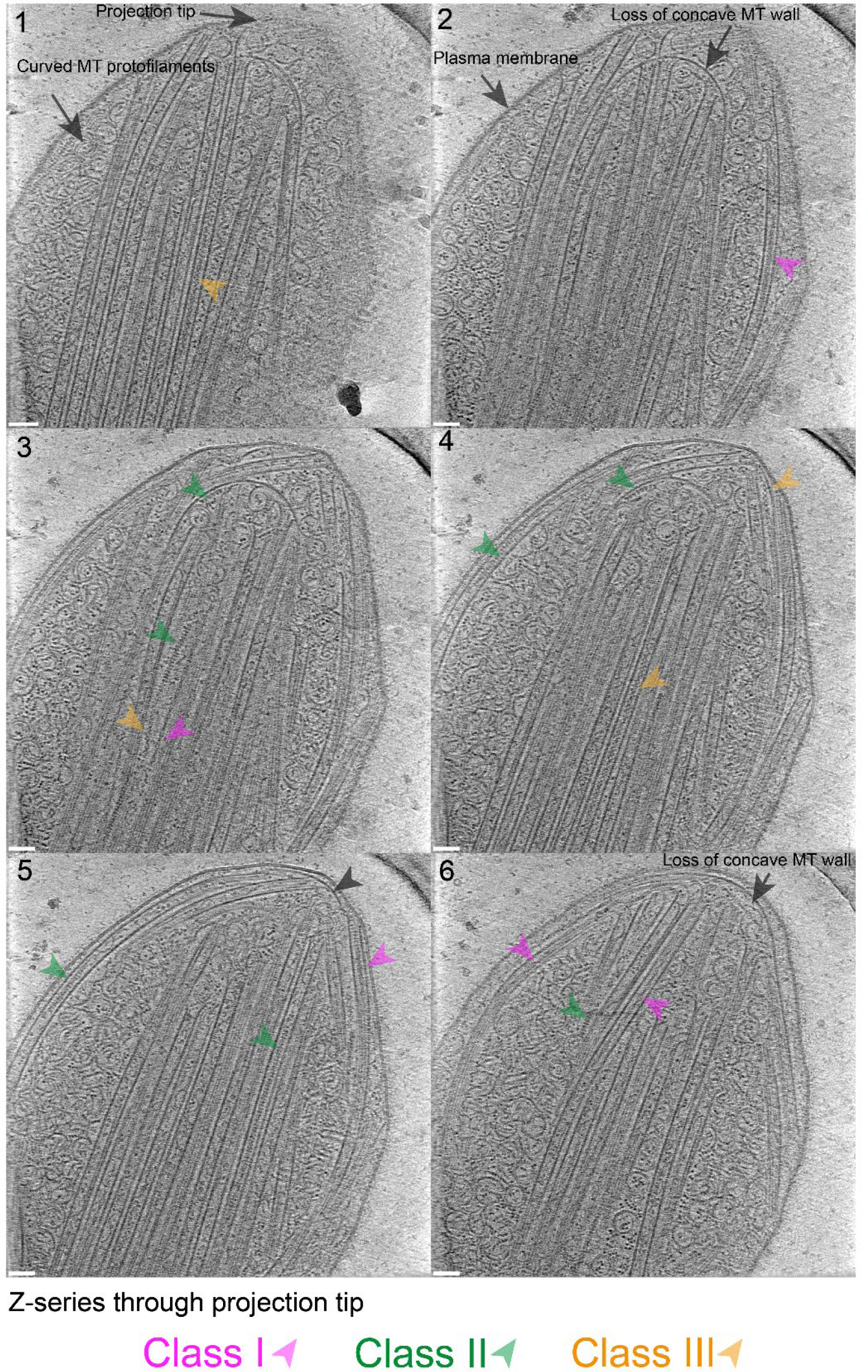
Microtubules are under structural and mechanical stress when they are extruded from HAP1 cells. Tomogram slices showing a series of images along the Z-axis of the tip of a projection. The three filament classes are highlighted by coloured arrows. A video showing this Z-series is provided in Supplementary Movie 1. Scale bar = 50 nm.

## Supplementary Movies

**Supplementary movie 1**. Image series through the tomogram depicted in **Fig. S2** showing a projection tip. Scale bar is 50 nm.

**Supplementary movie 2**. Image series through the tomogram depicted in **Fig. 3C** showing microtubules in a FIB-milled platelet. Scale bar is 50 nm.

## Material and Methods

### HAP1 cell culture and fluorescence imaging

HAP1 cells were grown in Iscove-modified Dulbecco Media with 10% FBS and penicillin/streptomycin at 37 °C in a 5% CO2 incubator. For fluorescence imaging, cells were prepared by an established protocol in Paul et al.[4] and stained using TUB 2.1 monoclonal antibody (1:1000, T4026, Sigma-Aldrich) detected with anti-mouse secondary antibody directly conjugated to Alexa Fluor 568 (1:500, A11004, Thermo Fisher Scientific), and stained for β-actin using 13E5 rabbit monoclonal antibody directly conjugated to Alexa Fluor 488 (1:500, 8844, Cell Signalling Technology).

### HAP1 cell culture and cryoET

HAP1 cells were grown as above. For kinesore treatment, Quantifoil R1.2/1.3 gold 300 mesh grids (Agar Scientific) were coated with 1mg/ml of fibronectin in a 6 well plate overnight at 37 °C. Cells were then plated at 0.5x10^5^ cells per well and incubated for 2 days at 37 °C. Cells were washed with Ringer’s buffer (pH 6.8) and treated with kinesore (Chembridge Corporation) at 0.2% concentration in DMSO (0.2% DMSO for control), in Ringer’s buffer (pH 6.8) for 1 hour, in a non-CO2 incubator. Grids were lifted out of the media and 10 nm gold fiducial markers were applied on the grids. Samples were blotted and plunge frozen in liquid ethane using a Leica EM GP plunge freezer. Cryo-EM grids were clipped and screened on Tecnai20 LaB6 TEM (FEI) at 200kV with a Gatan 626 cryo-transfer holder and sent to Diamond Light Source Electron Bio-Imaging Centre (eBIC) for tomography data collection at 64,000x magnification on a Titan Krios microscope with Falcon4 detector and a 5eV slit (Thermo Fisher). Tomographic series were acquired using a dose symmetric scheme, with increments of 3 degrees. Sample shown in Fig. S2 of a projection tip were acquired as part of the dataset reported in [4] and processed as described there.

### Tomographic reconstruction of HAP1 projections

The IMOD packages newstack and alignframes were used to order and motion correct the tilt series as part of a custom Bash script (Dr Mathew McLaren, University of Exeter), and reconstructed into a tomogram using weighted back projection with 5 Simultaneous Iterative Reconstruction Technique like filter on Etomo (IMOD). Reconstructions were performed using IMOD and its Etomo interface [29]. The tomogram presented in Fig. S2 was acquired as part of the dataset reported in [4] and processed as described there.

### Layer line analysis and helical reconstruction

Fourier transforms of 2D projection of extracted filament volumes were performed and layer line positions were measured using Fiji (ImageJ). Filaments of interest were extracted from 3dmod to ImageJ to straighten the filaments and invert densities. Real space helical reconstruction was performed using dimensions (axial rise 27.5 Å and subunit rotation 162°) calculated from the layer line analysis. Cofilin-actin (PDB - 3J0S) and actin (PDB - 8D17) were docked into helical reconstruction models in UCSF Chimera.

### *In vitro* reconstitution and cryoET

Tubulin (Cytoskeleton Inc: HTS03-A) and actin (Cytoskeleton Inc: APHL99) were co-polymerised at 4 mg/ml and 0.4 mg/ml respectively in a modified TicTac buffer (10 mM HEPES, 80 mM PIPES (pH 6.8), 50 mM KCl, 5 mM MgCl_2_, 1 mM EGTA, 1 mM GTP, 2.7 mM ATP, 1 mM DTT) [16] at 37°C for 1 hour. Cofilin (Cytoskeleton Inc: CF01-A) was added to the mixture before polymerisation at 0.2 mg/ml. 5 μl of the polymerised mixture was pipetted onto Quantifoil R1.2/1.3 copper 300 mesh grids (Agar Scientific) and plunge frozen in liquid ethane using a Leica EM GP plunge freezer. CryoEM grids were clipped and imaged on Talos Arctica (FEI) at 63,000x and a dose symmetric scheme with increments of 3 degrees from -60 to 60 degrees and reconstructed as above.

### Platelets isolation, FIB-milling, and cryoET

Platelets were isolated as described previously [30]. Briefly blood from healthy drug-free volunteers was drawn into 3.2% (w/v) trisodium citrate according to local NHS research ethics approval (20/SC/0222) and the Declaration of Helsinki. Blood was centrifuged at 1000 rpm for 17 min at room temperature (RCF = 180g) and platelet-rich plasma (PRP) was collected and supplemented with acidified Acid Citrate Dextrose (ACD) 1/7 (v/v) and apyrase (0.02 U/ml). Platelets were subsequently pelleted (1700 rpm/10 min) and washed with CGS (13mM trisodium citrate, 30mM glucose, 120mM sodium chloride) supplemented with 0.02 U/ml apyrase, before resuspension in modified Hepes-Tyrodes buffer (145 mM NaCl, 1mM MgCl_2_, 3 mM KCl, 10 mM HEPES pH 7.3) supplemented with 5.5 mM D-glucose and 0.02 U/ml apyrase, at a concentration of 4x10^8^ platelets/ml. Quantifoil R1.2/1.3 gold 300 mesh grids (Agar Scientific) were coated with Collagen Related Peptide at 50 μg/ml overnight at room temperature and blotted off or left uncoated. We did not observe a difference between uncoated and coated samples, and so data presented are pooled sets. 5 μl of isolated platelets were pipetted onto the grids and left for 5 minutes, blotted and plunge frozen in liquid ethane using a Leica EM GP plunge freezer. CryoEM grids were clipped and screened on a Talos Arctica (FEI).

Focused Ion Beam milling (FIB-milling) was performed at eBIC on an Aquilos cryoFIB/SEM (Thermo Fisher) to produce lamellae at 12 degree tilts. The samples were then imaged using a Titan Krios with a Falcon 4i detector at 300 keV at 53,000x with a dose symmetric scheme from -45 degrees to 69 degrees. The tomograms were reconstructed on the eBIC processing pipeline using AreTomo [31].

